# Sub-cellular Imaging of the Entire Protein-Coding Human Transcriptome (18933-plex) on FFPE Tissue Using Spatial Molecular Imaging

**DOI:** 10.1101/2024.11.27.625536

**Authors:** Rustem Khafizov, Erin Piazza, Yi Cui, Michael Patrick, Evelyn Metzger, Daniel McGuire, Dwayne Dunaway, Patrick Danaher, Margaret L Hoang, Andrew Grootsky, Megan Vandenberg, Shanshan He, Rachel Liu, Michael McKean, Michael Rhodes, Joseph M Beechem

## Abstract

Single-cell RNA-seq revolutionized single-cell biology, by providing a complete whole transcriptome view of individual cells. Regrettably, this was accomplished only for individual, tissue-dissociated cells. High-plex spatial biology has begun to recover the x, y, and z-coordinates of single-cells, but typically at the expense of far less than whole transcriptome coverage. To solve this problem, Bruker Spatial Biology has accomplished a commercial-grade panel (CosMx^®^ Spatial Molecular Imager Whole Transcriptome Panel (WTx)), using 37,872 imaging barcodes, capable of sub-cellular imaging of the entire human protein-coding transcriptome. The imaging barcodes are encoded with 156 bits of information (4 on-cycles and 35 dark-cycles per code), at a Hamming Distance of 4 from each other to achieve a very low false-code detection. Key to achieving this high-plex capability was the ability to manufacture imaging barcodes that require no in-tissue amplification (every barcode is manufactured under GMP to contain exactly 30 fluorescent dyes) and uniform, size-exclusion purified, extremely small imaging barcodes (∼ 20 nm). A detailed study of six different human FFPE tissue types was performed (Colon, Pancreas, Hippocampus, Skin, Breast, Kidney), yielding over 5.4 billion transcripts from 2.7 million cells. We counted over 1,550 transcripts-per-cell on average and observed 900 unique genes per cell (measured as the median). Single fixed-cells containing well over 10,000 subcellularly imaged transcripts were accomplished. Advancing single-cell imaging to the whole transcriptome level opens a single unified approach to accomplish essentially all single-cell experiments, both imaging and non-imaging. Depending upon the sample type (e.g. fixed-cells, organoids, tissue sections, etc.), the transcripts per cell and genes per cell measured using the whole transcriptome panel often exceeds that obtained by the highest-resolution single-cell RNA-seq, can be performed on a single 5 µm FFPE tissue section, with no dissociation bias (every cell is counted). Pathway analysis within the tumor bed of a colon adenocarcinoma sample found evidence of enrichment in pathways suggestive of an aggressive tumor type, and localized ligand-receptor analysis showed spatially restricted patterns related to adhesion, migration, and proliferation. The high-dimensional whole transcriptome data is streamed directly to a cloud-based Spatial Informatics Platform, allowing for the scalable processing of millions-of-single-cells and billions-of-transcripts per operation. The WTx data are combined with high-resolution antibody-based cell-morphology imaging and data-driven machine-learning cell segmentation algorithms, to generate the most complete view of single cell and sub-cellular spatial biology that has ever been obtained.

## Introduction

The shift from bulk sample analysis to single-cell analysis has revolutionized our understanding of biological systems, providing unparalleled insights into tissue organization, pathophysiological changes, and treatment responses. Traditionally, bulk transcriptomic techniques averaged the gene expression signals across thousands or even millions of cells, masking critical heterogeneity within tissues. The advent of single-cell technologies, however, has been transformative, enabling researchers to deconstruct this cellular complexity and observe how individual cells contribute to both normal physiology and disease pathogenesis. Over the last decade, single-cell RNA analysis has made tremendous strides, catalyzing breakthroughs in projects including the Human Cell Atlas, BRAIN Initiative, Cancer Moonshot, and next-generation therapeutics (*e.g.,* CRISPR medicines and mRNA vaccines)^1–5^.

Single-cell RNA analysis can be broadly classified into two main technical approaches: sequencing-based and imaging-based methods. Both approaches offer unique strengths while also facing inherent limitations.

Single-cell RNA-sequencing (scRNA-seq) has become the cornerstone for transcriptomic research due to its ability to provide comprehensive gene expression profiles from individual cells. By capturing and sequencing the entire transcriptome from each cell, scRNA-seq offers a deep and unbiased view of cellular states. Initial RNA-seq experiments were on just a handful of cells (a few hundred to thousand)^6^. The initial challenge was preparing individual cells, but the development of encapsulation methods led to a massive increase in cellular throughput^7^. However, this approach comes with significant drawbacks: the requirement to dissociate tissues into single-cell suspensions introduces biases by disrupting cellular environments and losing certain rare, fragile or densely packed cell types^8^. Additionally, scRNA-seq workflows are cost-intensive and suffer from data impurity, such as the formation of multiplets in droplet-based methods, where multiple cells are inadvertently captured in the same droplet, confounding the downstream analysis. Critically, the spatial information – the native tissue context and positional data of each transcript within a cell – is entirely lost in scRNA-seq, limiting its ability to correlate molecular information with cellular morphology or tissue architecture.

On the other hand, imaging-based RNA techniques trace their conceptional origins to *in situ* hybridization (ISH), a method that allows for the precise visualization of individual RNA molecules within cells^9^. Fluorescence ISH (FISH) offers remarkable subcellular resolution and provides nanometer-level localization accuracy. This makes imaging-based techniques ideal for studying the spatial distribution of transcripts in their native environments. However, these methods face significant multiplexing limitations. The number of distinct RNA species that can be detected simultaneously is constrained by spectral overlap, fluorophore availability, and signal-to-noise ratios within optically crowded environment. Over the past years, a variety of barcoding strategies were developed to achieve multiplexed FISH. The plex of initial FISH methods was 2 in isolated cells^10^, and the largest plex reported was 10,000 in mouse brain^11^. The growth of spatial imaging has been covered in several reviews^12^. However, imaging-based approaches have not matched single-cell sequencing by profiling the entire transcriptome in a single experiment.

To address the limitations inherent in these single-cell techniques, recent innovations have given rise to *in situ* capture and deterministic barcoding-based spatial transcriptomics tools. These methods aim to preserve spatial context while enabling high-throughput RNA profiling. Although spatial transcriptomics provides an intermediate solution by allowing whole-transcriptome coverage with preserved positional information, these approaches cannot localize individual transcripts or delineate single-cell boundaries, leaving a gap in our ability to integrate comprehensive transcriptomic and spatial data at the single-cell level^13,14^.

In response to these challenges, 37,872 imaging barcodes were developed to probe 18,936 human protein coding transcripts (the CosMx Whole Transcriptome (WTx) assay), enabling comprehensive whole-transcriptome profiling while maintaining spatial sub-cellular resolution down to the level of individual transcripts within cells. The barcodes are cyclically imaged *in situ* on the CosMx Spatial Molecular Imager^15^ (SMI), overcoming the loss of spatial information observed with scRNA-seq, while dramatically expanding the multiplexing capacity compared to traditional imaging techniques.

This assay is validated across a variety of sample types, including highly challenging FFPE tissues, and has the potential to reshape the landscape of single-cell analysis by delivering both transcriptomic depth and spatial context at unprecedented throughput, cost-effectiveness, and resolution. Researchers can now achieve a more complete understanding of cellular behavior within intact tissue environments, opening new avenues for biological discovery and therapeutic intervention.

## Results

### CosMx SMI Whole Transcriptome Design

The CosMx Human WTx assay consists of *in situ* hybridization (ISH) probes designed to target all protein-coding genes of the human transcriptome. We designed 37,890 ISH probes targeting 19,867 human genes covering >99.5% of the annotated protein-coding genes curated by the HUGO Gene Nomenclature Committee (HGNC^15^) for which there was available sequence in NCBI RefSeq^16^. We did not fail to design ISH probes for all protein-coding genes, and designed specific ISH probes to even very short transcripts (**Figure 1A**). Probes were aligned to the T2T genome and their relative locations along each chromosome are shown in **Figure 1B**. Two ISH probes were designed for each target gene, each designed against a different portion of the transcript. The two ISH probes for a single target are also referred to as “tiles”, as they “tile” a target transcript. Given high homology within some gene families, a subset of the targets hit multiple genes. In these cases, targets that hit the same set of genes were removed to prevent redundancy. Mitochondrial and nuclearly-encoded constitutive highly expressing genes (10 total) were not put into the panel, but can be added as a custom “add-on” to the panel if desired (see Methods). We additionally designed 100 negative control ISH probes, present as 50 negative control targets, against synthetic sequences from the External RNA Controls Consortium (ERCC) set^17^. The ERCC sequences have the same properties as mammalian sequences but without similarity to any known transcripts. All target genes included in the panel are listed in Supplementary Table 2.

**Figure 1.**
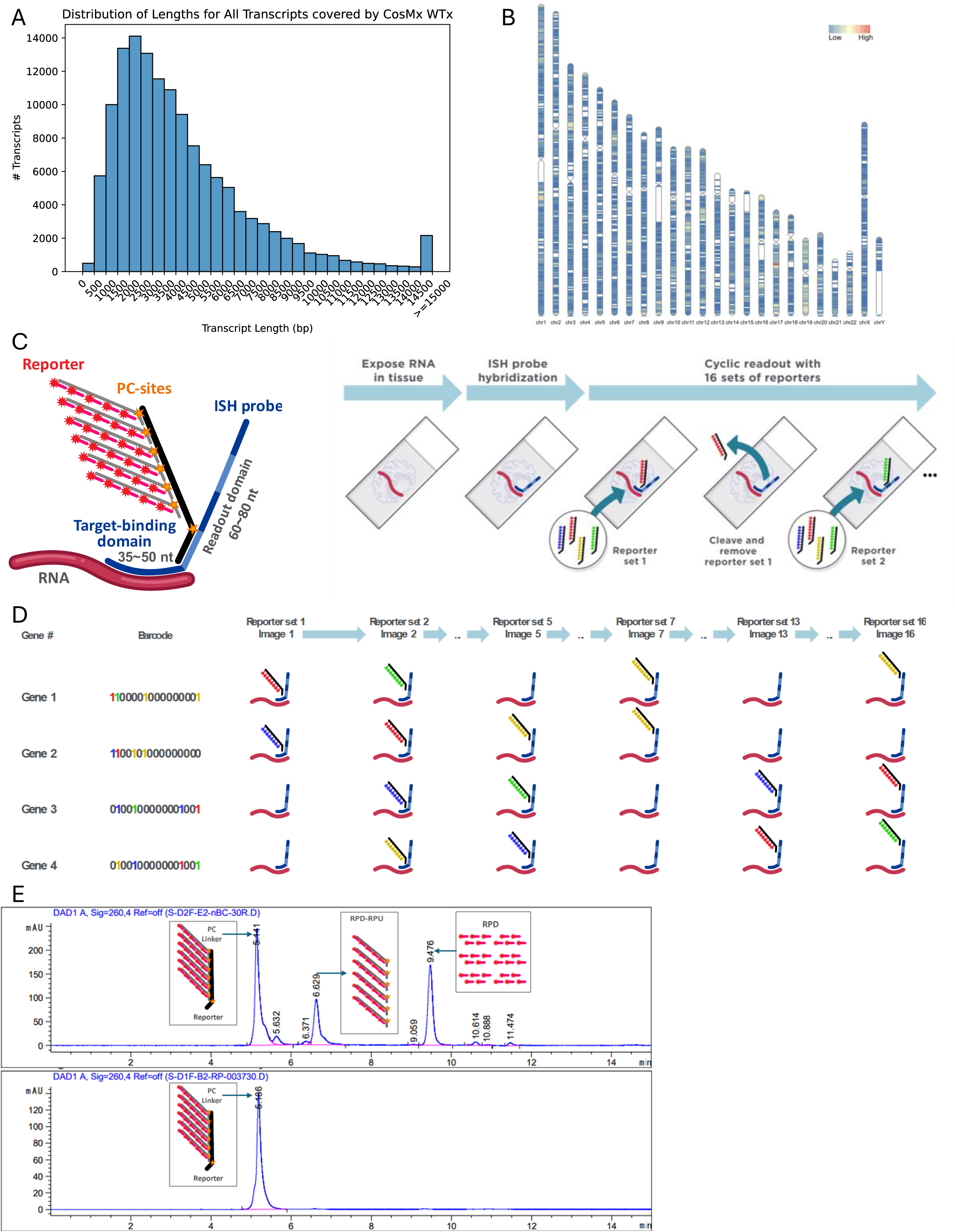
Design of the CosMx Whole Transcriptome Assay**. (A)** Distribution of sizes of all transcripts targeted by WTx probes is shown as a histogram. All transcripts greater than or equal to 15 kb in length are binned. **(B)** Gene density plot overlayed on the T2T genome karyotype demonstrates density of probe target regions across the chromosomes. **(C)** Schematic description of the ISH probe (blue), reporter design and illustration of the RNA assay workflow. The ISH probe consists of the target-binding and readout domains, the former being a 35–50-nt DNA sequence that hybridizes with target RNA, and the latter can be hybridized with a unique reporter. Each Reporter is conjugated with one of four fluorophores, and will be detected as one of four colors (blue, green, yellow or red) in SMI images. Each reporter has a controlled number of 30 dyes with six photocleavable sites to efficiently quench signals by UV illumination and a washing step before each cyclic reporter readout. First, the FFPE slide undergoes standard tissue preparation to expose RNA targets for ISH probe hybridization. Then the sample is assembled into a flow cell and loaded onto an instrument for cyclic readout with 39 sets of reporters. Because each reporter set contains four reporters with four different fluorophores, 156 unique reporters are used in the SMI assay to bind to the different reporter-landing domains on ISH probes. Following each set of reporter hybridization, high-resolution Z-stacked images are acquired followed by cleavage and removal of fluorophores from the reporters before incubation with the next set of reporters. **(D)** Each target is assigned with a unique 39-digit barcode with 4 “on” spots (labeled as either B,G,Y or R) and 35 “off” spots (labeled as “0”). Each colored letter of the barcode indicates the presence of the reporter that is associated with the target in the specific reporter hybridization round and its color indicates the fluorophore of the hybridized reporter. “0” means that no reporter binds to the ISH probe in that hybridization round, and the target should be silenced or blank in that round of imaging. For each gene, 4 reporters will sequentially bind to the 4 designated reporter landing domains of the ISH probe throughout the 39 rounds of cyclic reporter readout. **(E)** Representative 260nm chromatograms for reporter assembly and purification are shown. Unpurified 30 dye reporters with Alexa Fluor-647 (top) shown in contrast to purified product (bottom). Each inset box represents what is represented by each chromatogram peak. Each element is hybridized in excess to ensure fully assembled reporters.

ISH RNA-targeting domains are 35–50 nucleotides in length, and were selected based on an iterative design process that considers thermodynamic profile, splice isoform coverage, potential for cross-hybridization with other transcripts, and potential for intramolecular interactions between probes (see Methods). Probes were synthesized individually and pooled.

### Readout and Reporter Chemistry

Each ISH probe can be divided into two functional domains: an RNA-targeting domain and a readout domain for the imaging-barcode oligos (**Figure 1C**).

The readout domain consists of four consecutive sequences that allow four individual SMI imaging barcodes (reporters) to bind sequentially. Each tile has its unique sequence in the target-binding domain, whereas both tiles targeting the same target share the same sequence in the imaging readout domain. The imaging readout domain of any single tile can uniquely identify the target gene (as small as 35 to 50 nucleotides), enabling high detection sensitivity in FFPE tissue, where RNA molecules may be highly fragmented. Each imaging reporter construct contains exactly 30 fluorescent dyes, with the fluorophore-conjugated oligos coupled to photocleavable (PC) linkers (**Figure 1C**). All reporters are single color, containing one of four fluorophores: Alexa Fluor-488, ATTO 532, Dyomics-605 or Alexa Fluor-647. The key advantages of the SMI reporter chemistry are high signal-to-noise ratio (SNR) detection over background fluorescence for accurate spot calling and fast fluorescent signal darkening, enabled by ultraviolet (UV) cleavage of dyes from the imaging barcodes. Manufacturing of the 30 dye reporter construct is tightly controlled and purified via size-exclusion HPLC (**Figure 1E**; Good Manufacturing Practices, ISO 9001). This approach ensures that the *in situ* imaging of every measured transcript has identical spot-size and fluorescence intensity, enabling this approach to scale to whole transcriptome levels without reagent-based detection bias. *In situ* enzyme amplification-based RNA transcript detection, such as rolling-circle-amplification, results in an uneven distribution of both spot-size and fluorescent intensity for RNA transcripts^18^.

For sample preparation, SMI utilizes standard FFPE ISH tissue processing methods to expose RNA targets, followed by the introduction of fluorescent bead-based 4-color fiducials (Bangs Labs, Fishers IN, custom product) that are fixed to the tissue to provide an optical reference point for cyclic image acquisition and multi-color registration. Following tissue processing and hybridization of ISH probes, slides are washed, assembled into a flow cell and placed within a fluidic manifold on the SMI instrument for RNA readout and morphological imaging. In the WTx RNA assay, the tissue is hybridized with 39 sets of fluorescent reporters sequentially, each reporter set containing four single-color reporter pools. The reporters specifically bind to ISH probes during the 39 rounds of reporter hybridization according to the barcode assigned to each gene (**Figure 1D**). After incubation of each set of reporters, high-resolution Z-stacked images are acquired for downstream analysis. Before incubation with the next set of reporters, photocleavable (PC) linkers are cleaved by UV illumination and released fluorophores are removed by washing. The SMI encoding scheme is designed to assign a unique barcode to each target transcript from a set of 156-bit barcodes (four color reporters in each readout round over 39 readout rounds), with Hamming distance 4 (HD4) and Hamming weight 4 (HW4). Every barcode is separated by an HD of at least four from all other barcodes to maximally suppress RNA decoding error. Every barcode has a constant HW4, in which each target is “on” (for potential reporter barcode binding) in four rounds and “off” in 35 rounds during the 39 rounds of reporter hybridization. It should be emphasized that even though a potential barcode set is “on”, does not mean that a reporter barcode will bind. The SMI-chemistry has been tuned, such that approximately Poisson-type-sampling occurs on the ISH-hybridized probes. This Poisson-sampled “on” and “off” signal allows for a facile expansion to a whole transcriptome scale (∼19,000), because only a subset of RNAs is “on” in any given cycle. Note that the nature of these on–off imaging signals represents a “deterministic super-resolution imaging system”, and hence each SMI imaging barcode can be located well below the diffraction limit of the imaging system. This super-resolution imaging generates a “location-error” for each transcript below 50 nm in both x and y coordinates (**Supplementary Figure 1**).

Similar to previously described^19^, the instrument takes image stacks in each field-of-view of 0.509 mm X 0.509 mm (n = 1 to 1149, 1149 to the maximum needed for imaging ∼ 3 cm^2^) after each reporter cycle. Primary analysis reduces these images down to an aligned list of spots in each color channel and cycle. Spots are sub-pixel localized by fitting with a weighted centroid, which was an improvement over the 2D paraboloid previously used^15^. The on-instrument detection platform has 10 CPU cores and supports up to 20 simultaneous processing threads, as well as a GPU card (NVIDIA RTX A4000, Santa Clara, CA), to detect spots in real time and power real-time cell segmentation analysis.

### Secondary Analysis

Given the dataset of reporter-binding events (RBEs) corresponding to spot location and imaging-barcode color, a decoding algorithm is used to infer location and target gene-identity of RNA transcripts. After candidate transcripts are identified as matching at least 3 of the 4 on-reporters in the gene’s barcode within a fixed 1.1 pixel neighborhood radius, machine learning models are used to infer quality scores for each transcript, based on ability to distinguish the decoded transcript from system control barcodes and negative probe controls (**Figure 2A-C**). These quality scores are used to infer a set of high-quality decoded transcript calls prior to tertiary analysis (**Figure 2D**). The machine learning models rely on features derived from the local neighborhood of each transcript, including a score of local barcode crowdedness and the spatial spread of observed RBEs (see Methods).

**Figure 2.**
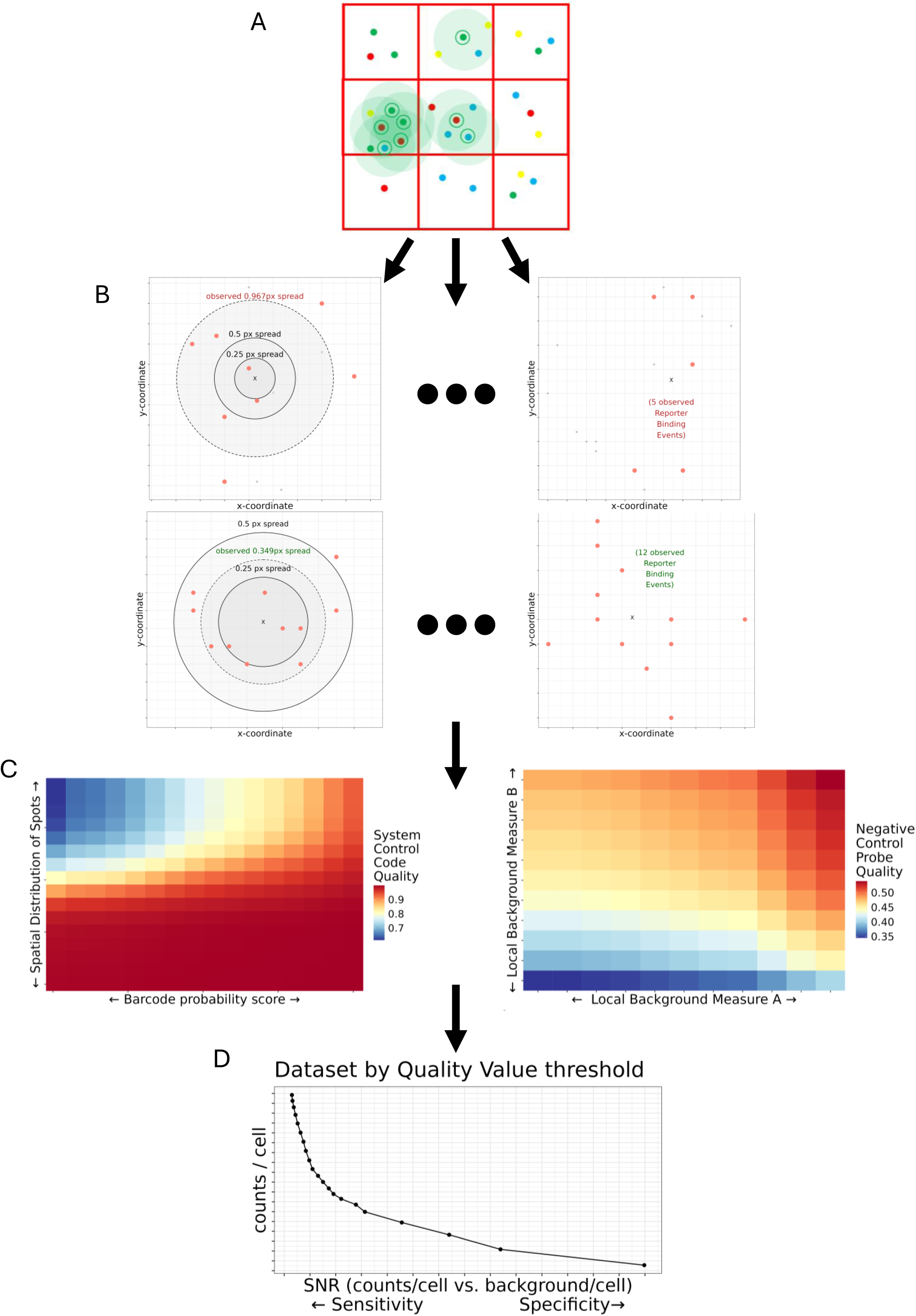
Overview of adaptive decoding algorithm. **(A)** Targets with at least 3 of 4 barcode spots within a search radius are identified, and **(B)** multiple metrics related to target call quality are computed, including the spread of observed reporter binding events, total number of reporter binding events, posterior probability score, and whether we observe a full (4-spot) or partial (3-spot) barcode. **(C)** These metrics are used to train two machine learning models, each predicting either SystemControl / Negative Control Probe as a binary outcome, and imposing common-sense monotonic constraints on each feature. **(D)** Overall quality value for each transcript is computed as combination of SystemControl and NegativeControl Probe quality scores. Transcripts are ranked by their quality value, and target calls below a specified quality threshold are discarded. Different quality value cutoffs represent a smooth tradeoff between sensitivity (transcripts/cell) and specificity (transcripts to background ratio).

### Performance of the WTx Assay on Cell Pellets

To measure WTx assay performance, we created an FFPE human cell pellet array from 36 cell lines (CPA36) previously characterized by bulk RNA-seq^20^. The 36 cell lines originated from 18 different tissue types and 17,072 of 18,935 (90.2%) genes are expressed at TPM >= 1 in at least one cell line in the Cancer Cell Line Encyclopedia (CCLE) database^20^. Three independent WTx experiments performed on the CPA36 demonstrated high reproducibility (R = 0.97 +/- 0.010) and five order-of-magnitude dynamic range of transcript counts (**Figure 3A**). CCLE bulk RNA-seq TPM values varied over four orders of magnitude, as expected from the deep sequencing per cell line (>100M reads^20^). We cross validated WTx single-cell counts by comparing CosMx WTx pseudo-bulk to CCLE bulk RNA-seq TPM. This comparison produced a “hockey stick” trendline (**Figure 3B**), consistent with previous studies^19^ and independent of detection method and platform^11,21^. Breakpoints occurred at TPM = 1.08 +/- 0.39 (**Figure 3C**), similar to previous data using the 1000-plex assay on SMI^19^. Above the breakpoint, WTx showed high correlation to bulk RNA-seq (R = 0.76 +/- 0.02, **Figure 3D** and see Figure 3B). Using bulk RNA-seq as ground truth (see Methods), we calculated sensitivity at 67.17 +/- 5.21% and specificity at 96.53 +/- 0.37%, comparable to published measurements for 1000-plex panel^19^. Lastly, we created a 6,000-plex (6K-plex) “nested” subpools of the WTx panel to test if increasing plex to whole transcriptome maintains quantitation of each individual gene. Indeed, high correlations were observed in the 6K-plex subpools compared to the same genes within the full ∼19,000-plex WTx panel (R > 0.98, **Figure 3E**), indicating that expression measurements were consistent across plex level. Based on these findings, we conclude that the CosMx platform provides reproducible measurement of gene expression scalable to the whole transcriptome.

**Figure 3.**
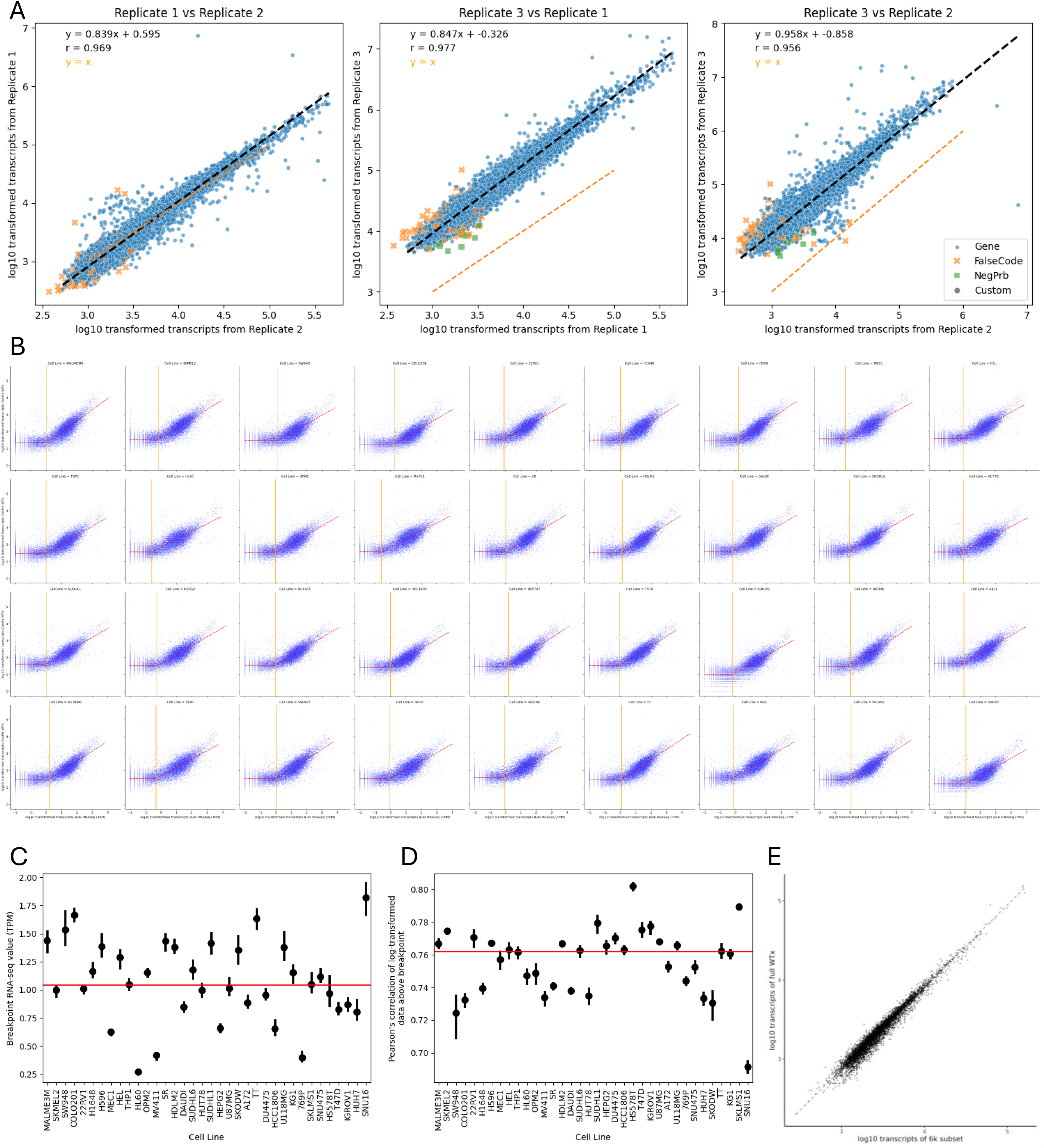
WTx assay performance in cell pellet arrays. **(A)** Three replicate CosMx runs are plotted by total sum of all transcript calls transformed to log10 scale. Replicate 1 and 2 are done at a small imaging scale (37 FOVs), while Replicate 3 is done at a large imaging scale (370 FOVs), suggesting that consistency is not compromised on slides with more FOVs. Blue dots denote transcript calls, orange “X”s denote system controls, and green squares denote negative controls. The black line shows the best fit line to transcript calls (omitting system and negative controls), while the orange line shows a line of slope = 1, and no intercept. **(B)** CosMx WTx RNA expression profiling is concordant with bulk RNA-seq. Log10 transformed CosMx data, the sum of all transcripts across a single FOV (y-axis), is compared against log10 transformed bulk RNA-seq data, in TPM, collected from the CCLE (x-axis) for 36 cell lines on a 5 µm thick section of FFPE cell pellet array. Red lines describe segmented regression, and orange lines show the log10 transformed estimated breakpoints from each cell line in TPM. **(C)** Estimated breakpoints for each cell line in TPM, across 12 FOVs (per cell line) and 3 replicate slides. Black lines show 95% confidence interval, and the red line shows median value. **(D)** Pearson’s correlation between log10 transformed RNA-seq and CosMx data, for all transcripts above the breakpoint, across 12 FOVs (per cell line) and 3 replicate slides. The black line shows 95% confidence interval, and the red line shows median value. **(E)** Correlation between the log10 transformed transcripts for the full WTx panel (y-axis) and the log10 transformed transcripts for a subset of 6064 targets (x-axis) that corresponds to the commercial 6k panel, with targets selected for further redesign temporarily removed. Black dotted line shows the best fit regression.

### Performance of the WTx Assay on Human FFPE Tissue Samples

To assess WTx performance on archival human tissue specimens, we profiled 5 µm FFPE sections from six different tissue types (colon, pancreas, skin, kidney, breast, brain) with an average area of 99.99 +/- 13.84 mm^2^ to generate 450,109.5 +/- 154,610 single cells with spatial context per tissue (**Table 1**). Neural and non-neural tissues were visualized on instrument with cell segmentation markers for nuclei, cell bodies, and plasma membranes (**Figure 4A**). Segmented cells showed a range of cell areas within and between tissue types (**Figure 4B**). For example, average cell area in pancreas was 71.99 µm^2^ compared to hippocampus at 167.76 µm^2^, consistent with expected cell sizes and composition within each tissue^22^. WTx profiling of the six tissue types revealed between 8,392-15,960 genes detected. 6,647 genes were detected in common across all tissues and between 106 - 843 were uniquely detected in one of the six tissue types (**Figure 4C**). Transcripts detected per cell was a median of 1,556 (965-2,248 range across tissues) and unique genes detected per cell was a median of 905 (627-1,252 range across tissues) (**Figure 4D**). We further assessed transcript assignment by analyzing the coincident detection of two mutually exclusive marker genes. For example, in colon cancer cells *IGKC* (plasma cells) and *EPCAM* (epithelial cells) were co-detected in 1.01% of cells while in hippocampus *SLC17A7* (excitatory neurons) and *C1QC* (microglia) were co-detected in 0.67% of cells (**Figure 4E**), suggesting valid cell segmentation within a diversity of tissue architectures. In summary, WTx captures tissue expression of large areas (1 cm x 1 cm) at cellular resolution across a range of archival human FFPE tissues.

**Figure 4.**
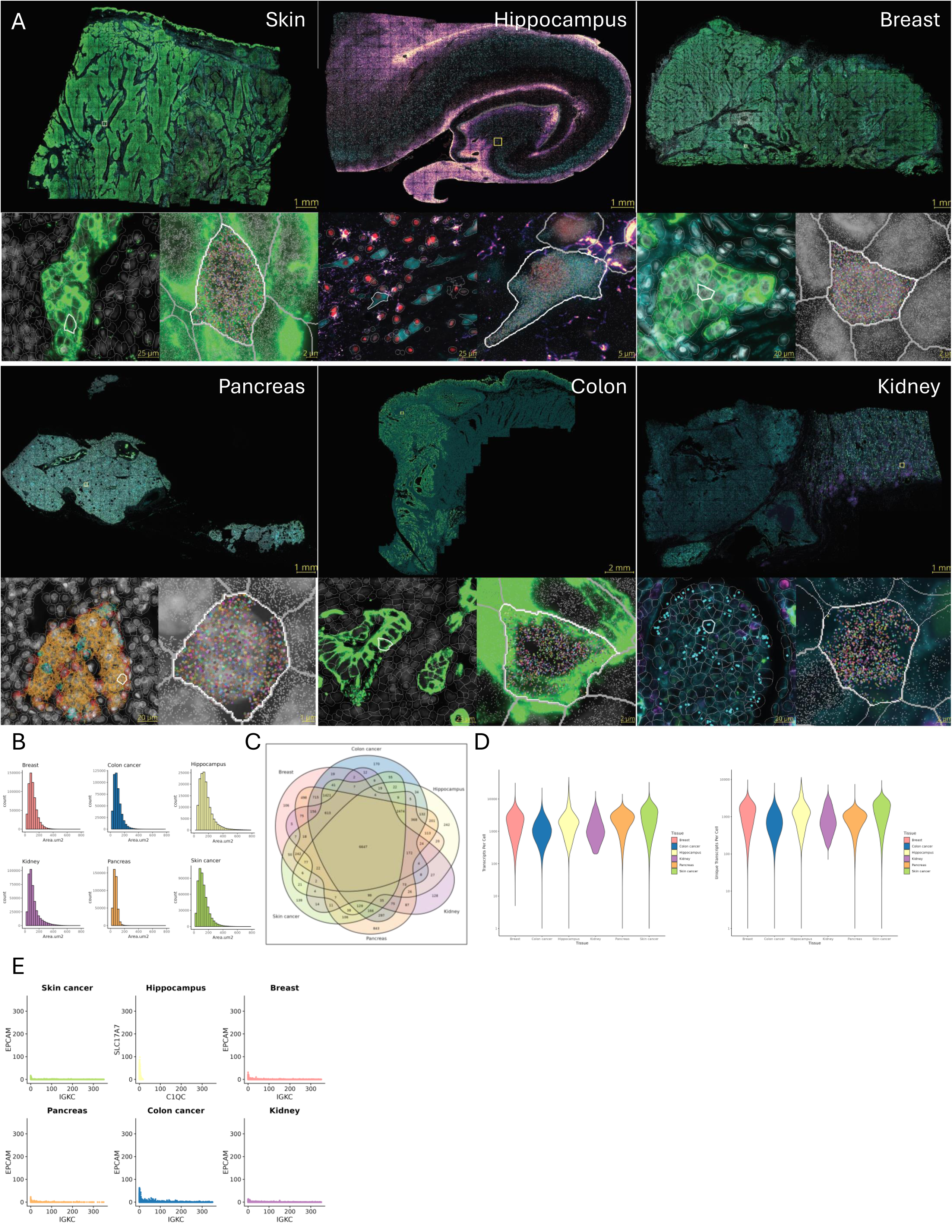
WTx assay performance in FFPE tissues. **(A)** Whole transcriptome spatial data segmented in six different tissues: Skin, Hippocampus, Breast, Pancreas, Colon, and Kidney. For each tissue, yellow squares in full image show the extent of the corresponding region-level (lower-left) inset and cell-level inset is shown in the lower right. In the cell-level insets, individual transcripts for the focal cell (white border) is shown with color and transcripts of non-focal cells appear as gray dots. Note that in some of the cell-level insets, individual fluorescence channels were removed for clarity. Within the region-level insets, white (focal cell) and gray (all other cells) polygons denote cell segmentation boundaries. Cell type maker genes are highlighted to show micro-tissue structure in Pancreas (red = *GCG* [alpha cells], orange = *INS* [beta cells], cyan = *SST* [delta cells]) and kidney (cyan = *VEGFA* [podocytes]). Immunofluorescent (IF) markers for all tissues except Hippocampus: green = PanCK, cyan = Membrane, gray = DAPI. IF markers for Hippocampus: red = histone, “magma” = GFAP, cyan = rRNA, gray = DAPI. In addition, magenta in Kidney = CD45. **(B)** Distribution of cell area in µm2 across diverse tissue samples. **(C)** 6-way Venn diagram describing the number of overlapping genes above the limit of detection in diverse tissue samples. **(D)** Distribution of total transcripts per cell and unique transcripts per cell across diverse tissue types. Y-axis transformed to log10 scale. **(E)** Barnyard plots showcasing specificity of cell type specific markers. For non-neuronal tissues, *EPCAM* (epithelial cells) is plotted against *IGKC* (plasma cells). For neuronal tissues, *SLC17A7* (excitatory neurons) is plotted against *C1QC* (microglia).

**Table 1.** Metrics associated with WTx assay performance in 6 different FFPE tissues.

| Metric | Colon | Pancreas | Hippocampus | Skin | Breast | Kidney |
| --- | --- | --- | --- | --- | --- | --- |
| Number of FOVs | 400 | 267 | 360 | 400 | 400 | 393 |
| Total Tissue Area (mm<sup>2</sup>) | 104.9 | 70.0 | 94.4 | 104.9 | 104.9 | 103.0 |
| Mean Single Cell Size (μm<sup>2</sup>) | 102 | 72 | 168 | 132 | 111 | 120 |
| Number of Cells in Analysis | 493,929 | 413,946 | 154,610 | 556,134 | 645,251 | 436,787 |
| Total Transcripts Assigned to Cells | 642,080,290 | 936,778,485 | 352,838,087 | 1,608,408,183 | 1,270,179,251 | 642,882,039 |
| Mean Transcripts per Cell | 1300 | 2263 | 2290 | 3031 | 1969 | 1472 |
| Maximum Transcripts per Cell | 22067 | 14214 | 46985 | 32877 | 31236 | 17300 |
| Mean Unique Genes per Cell | 793 | 865 | 1371 | 1506 | 1190 | 912 |
| Mean Transcripts per μm<sup>2</sup> | 12.75 | 31.43 | 13.63 | 22.89 | 17.7 | 12.3 |
| Mean Negative counts per Cell | 0.729 | 0.625 | 1.383 | 0.908 | 0.785 | 0.510 |
| Mean Negative per Plex per Cell | 0.015 | 0.013 | 0.028 | 0.019 | 0.016 | 0.010 |
| Mean False Codes per Plex per Cell | 0.005 | 0.019 | 0.012 | 0.009 | 0.007 | 0.004 |
| Signal-to-Noise Ratio (global) | 4.71 | 9.55 | 4.36 | 8.56 | 6.62 | 7.62 |

### Benchmarking of Spatial Whole Transcriptomes against scRNA-seq

To benchmark the performance of the CosMx WTx assay against the widely used single-cell RNA sequencing (scRNA-seq) platform, we analyzed adjacent tissue sections from the same FFPE colorectal carcinoma block from a 10X certified service provider (Azenta Life Sciences, South San Francisco, CA). For the scRNA-seq data, two curls of 25 μm-thick sections were processed, yielding 156,250 dissociated cells (utilizing the Miltenyi FFPE tissue dissociation kit). Of these, 10,000 cells were subjected to RNA barcoding and profiling using the Chromium Flex assay, resulting in a final dataset with 6,257 cells, consistent with Chromium recovery rates of 60-70% from FFPE tissues^23^.

For the CosMx WTx assay, a single 5 μm tissue section from the same tissue block was imaged and analyzed, successfully segmenting 493,929 cells, with a total of 642 million total transcripts (**Figure 5A**). It is worth noting the large difference in total cell counts between the two methodologies: SMI resolves ∼ 0.5 million single cells per 5 µm tissue section, whereas the Chromium Flex assay resolves 60,000 to 70,000 cells per 5 µm of tissue input^23^.

**Figure 5.**
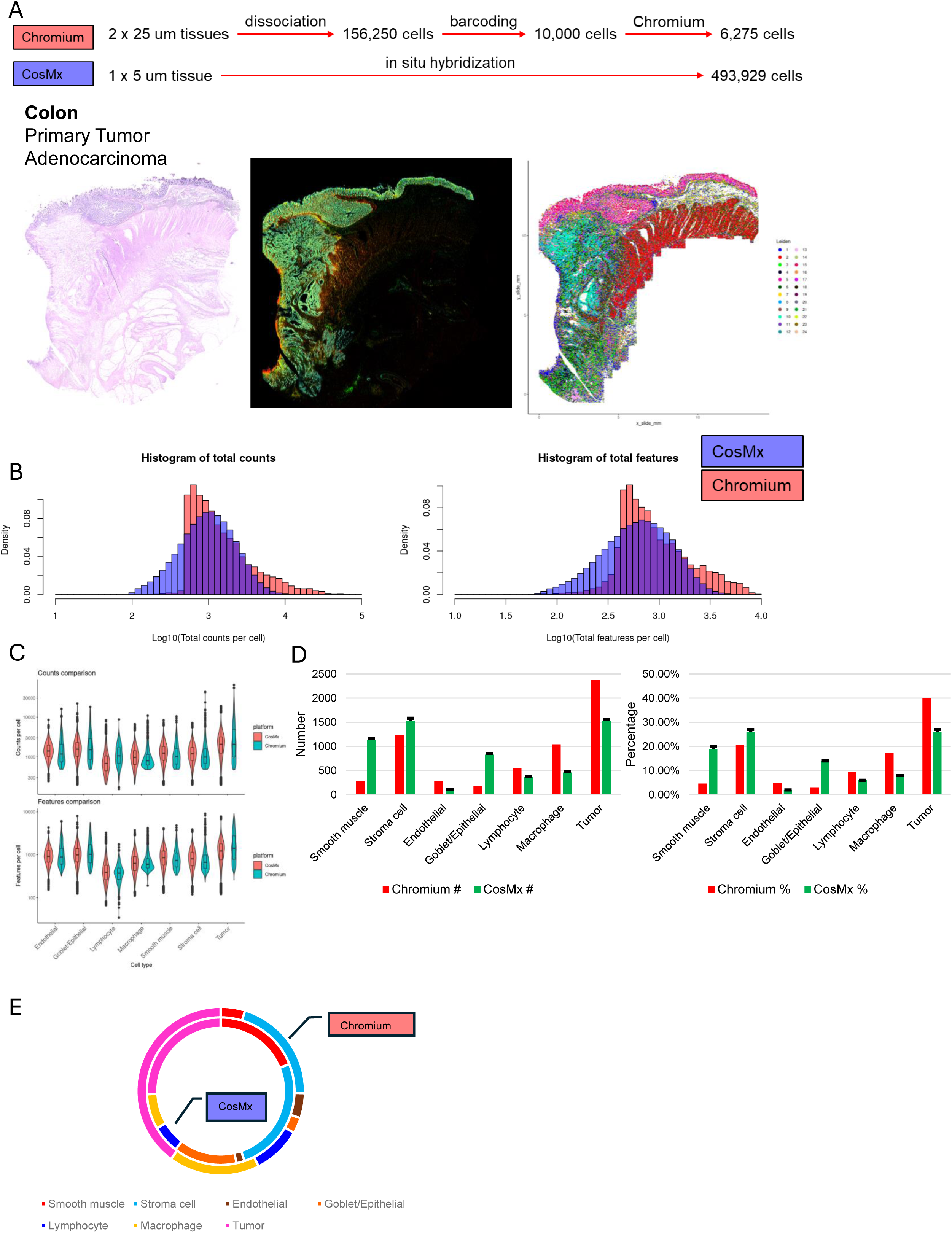
Performance comparison between CosMx WTx and single-cell RNA sequencing in FFPE colon cancer tissue. **(A)** Workflow schematic and throughput for Chromium FLEX and CosMx WTx assays. CosMx enables multi-modal integration of H&E, immunofluorescence and transcriptomics data on the same tissue section. **(B)** Histograms of detected total transcript counts per cell and gene features per cell from both platforms. **(C)** Cell type-specific comparison between the platforms (a coarse-level of 7 primary cell types was adopted). **(D)** Cell compositions of the tissue obtained from Chromium and CosMx were compared. To balance the difference in sample size, the CosMx dataset was subsampled: ∼6,000 cells were randomly chosen and mean composition was calculated (n = 100, mean + s.d.). **(E)** Cell type composition for entire dataset population from both platforms.

In terms of detected transcript counts and gene features, the CosMx WTx dataset exhibited a more uniform distribution compared to scRNA-seq (**Figure 5B**). This likely reflects the absence of multiplets – an issue that often arises in droplet-based scRNA-seq technique (especially for FFPE tissue), where multiple cells can be captured in a single droplet, confounding the analysis^24^. Both datasets underwent coarse cell clustering and annotation, identifying seven primary cell types. Across these cell types, CosMx WTx demonstrated highly comparable or improved detection efficiency relative to scRNA-seq (**Figure 5C**).

Notably, we observed differences in the cell composition between the datasets (**Figure 5D-E**). In the scRNA-seq data, certain cell populations, such as normal muscle and epithelial cells, were significantly underrepresented, despite these cell types constituting a large portion of the original tissue (**Supplementary Figure 2**). This discrepancy can be attributed to the challenges of dissociating densely packed or irregularly shaped cells during the scRNA-seq workflow^8^. By contrast, the CosMx WTx assay, free from dissociation-related biases, provided a more accurate representation of the tissue’s cellular composition (see detailed comparison metrics in **Supplementary Table 1**).

### Use of CosMx WTx on Cultured Cells

A specialized sample preparation protocol was developed for conducting CosMx WTx whole-transcriptome assays in cultured cells. Notably, the heat-induced epitope retrieval step was omitted, and the non-selective proteolysis with endoproteinase-K was substituted with a milder, selective proteolysis using endoproteinase GluC. As demonstrated in **Figure 6A**, the initial morphology staining effectively delineated cell boundaries, enabling precise segmentation. Importantly, the performance of the WTx assay in cultured cells frequently detected significantly more transcripts compared to FFPE tissues, likely due to superior RNA quality, lower intra-cellular cross-linking and high-contrast morphology staining. For example, in the highlighted Cell ID_232 (red arrow), a total of 10,586 transcripts from 4,844 genes were detected. Furthermore, the assay recorded both lateral and axial positional information for each detected transcript.

**Figure 6.**
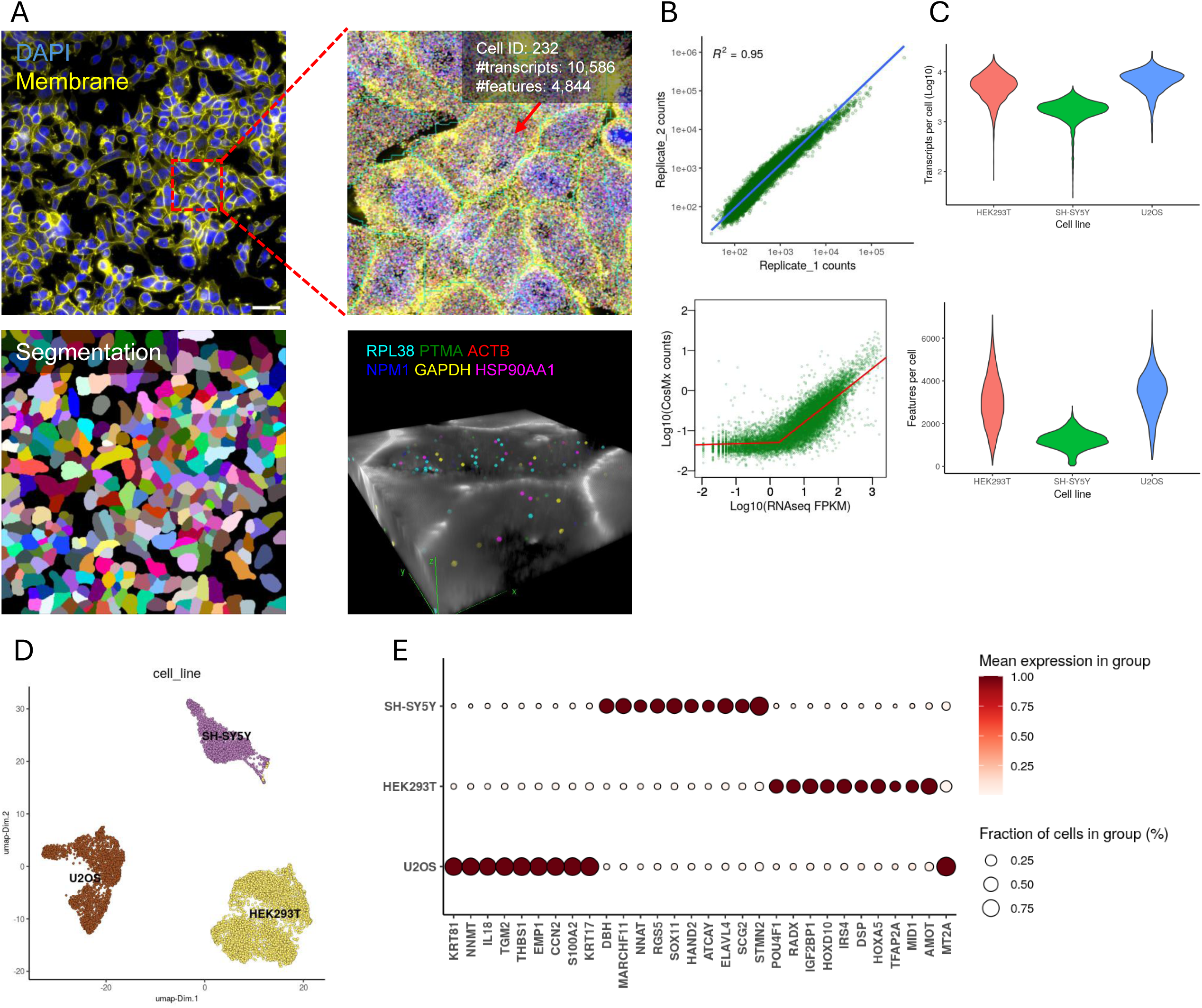
Performance of CosMx WTx in cultured cells. **(A)** Upfront morphology staining cell membrane (yellow) and nuclei stain (blue) shown in top left. In the image below, cell segmentation based on the above markers shown. Red dashed line in top left image delineates the boundaries of the area shown in the top right image. In the top right image, the red arrow points to one cell of interest (ID 232). Additional red box inset in the top right image shows area for bottom right image. Bottom right image shows 3D localization of transcripts in cells. **(B)** Assay reproducibility was validated in two independent runs using HEK293T cells, as shown in the upper correlation plot. Correlation with bulk RNA-seq data in the same line is presented in the lower plot. **(C)** Violin plots showing mean transcript counts and gene features per cell detected in 3 cell lines (n > 50,000 cells per cell line). **(D)** Upon dimension reduction, data for the three cell lines are presented via UMAP. **(E)** The top differentially expressed genes across the cell lines are shown with heatmap.

The CosMx WTx assays also exhibited high reproducibility in cultured cells and demonstrated a strong correlation with bulk RNA-seq data above the assay’s limit of detection threshold (**Figure 6B**). On average, CosMx WTx detected approximately 5,849 transcripts per cell in HEK293T (embryonic kidney epithelial), 1,866 transcripts per cell in SH-SY5Y (neuroblastoma), and 7,294 transcripts per cell in U2OS (osteosarcoma) cell lines (**Figure 6C**). The assay also supported robust cell clustering and typing following dimensionality reduction, with each cell line distinctly localized in UMAP space (**Figure 6D**). For each cell line, a highly specific transcriptomic signature facilitated unambiguous annotation (**Figure 6E**).

The CosMx WTx assay represents a significant increase in data density for subcellular transcript localization. In order to efficiently process multiple millions of cells and billions of transcripts, CosMx SMI streams data directly to a cloud-based AtoMx Spatial Informatics Platform (SIP). SIP is capable of the real-time automated scaling to the number of CPUs, GPUs, and memory required to analyze data of this magnitude. For example, for AtoMx Version 1.3, CPUs can scale to a maximum of 183, and memory can scale to 27 GB, allowing the facile processing of data sets that may approach 4 billion transcripts. AtoMx currently houses over 1.4 billion spatially resolved single cells, with over 2 petabytes of data storage. The initial datasets captured by CosMx for our tissue studies in Figure 4 ranged in size from 5.3-5.9 TB. The AtoMx SIP informatics suite includes a multi-modal, AI cell segmentation toolkit, iterative and custom analysis pipelines, and interactive image exploration. It significantly reduces processing times and addresses data scalability challenges encountered with open-source solutions including Squidpy^25^ and Seurat^26^.

### Application of the CosMx WTx Assay to Colon Cancer

To demonstrate the cellular resolution, sensitivity, and comprehensive spatial profiling capabilities of the WTx panel, 493,834 cells from an FFPE slide of a colon adenocarcinoma were analyzed. Following cell typing of the 414,005 cells that passed quality control (**Figure 7A**), locally restricted (**Figure 7B**) and global (**Figure 7C**) biological pathway changes were tested using the PROGENy database^27^ in the python package LIANA+^28^.

**Figure 7.**
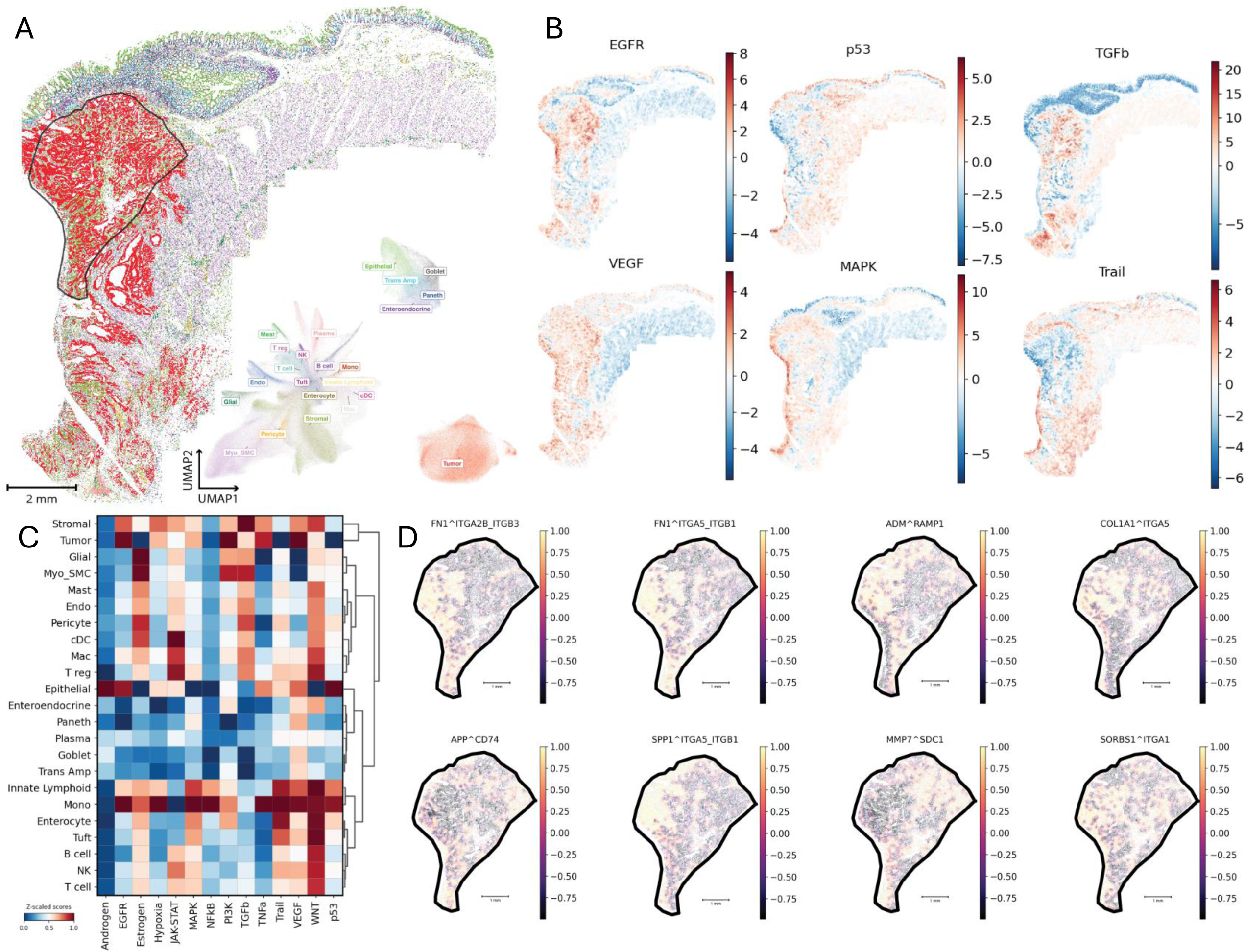
Pathway and Ligand-Receptor interactions identify global and spatial intratumor heterogeneity in colon cancer. **(A)** Major cell types identified using CosMx SMI WTx data for 414,005 cells. Cells are shown in their native XY space and UMAP space. Large tumor nest (black outline) is used for LR evaluation (panel D). **(B)** Local pathway enrichment scores for select PROGENy pathways. **(C)** Global view of pathway enrichment for all major cell type classifications. **(D)** Local LR scores for a given LR complex. Shown here are the eight complexes with the highest region-wide variability.

Within the tumor nest, an enrichment in EGFR, TGF-beta (TGFb), and VEGF pathways was observed, along with a reduction in the tumor necrosis factor-(TNF)-related apoptosis-inducing ligand (TRAIL) pathway, consistent with an aggressive tumor phenotype^29–32^. EGFR overexpression is associated with increased cell proliferation, invasion, and metastasis^29^. Increased expression of TGF-beta has been shown to enhance tumor cell survival and, together with VEGF, promote angiogenesis^30,31^. Finally, Gong et al.^32^ found that increased administration of TRAIL in mice promoted apoptosis in colon tumor cells. The lack of TRAIL pathway activity in the tumor further supports the conclusion that the tumor has evaded the immune response^33^. An enrichment of the MAPK pathway along the perimeter of the tissue also supports tumor aggression^34^, with monocytes, innate lymphoid cells, and stromal cells appearing dysregulated in addition to tumor cells (**Figure 7B**).

Given the larger size of the profiled tumor nest (10.1 mm^2^; 87,980 cells; 22% of cellular area), intratumor variation was investigated via local ligand-receptor (LR) analysis. Among the most locally variable, significant LR interactions, FN1 (ligand) was found to be interacting with integrin receptors ITGA2B/ITGB3 and ITGA5/ITGB1. ITGA5/ITGB1 also showed intratumor variance with SPP1 (osteopontin) (**Figure 7D**). These interactions suggest spatially restricted differences in cell adhesion, migration, and proliferation within the tumor^35–37^.

Taken together, the patterns observed suggest that different regions of the tumor may be characterized by distinct communication patterns and microenvironments.

## Discussion

The CosMx Whole-Transcriptome (WTx) assay represents a significant milestone in the field of spatial transcriptomics, achieving comprehensive whole-transcriptome coverage with sub-cellular resolution in an imaging-based format that requires no sequencing. Built on cyclic *in situ* hybridization chemistry, the CosMx WTx assay bridges critical gaps in the evolution of spatial and single-cell transcriptomic techniques. This study showcases how scalable, enzyme-free hybridization-based barcoding combined with wide-field optics, low volume microfluidics, antibody-based cell morphology markers, and adaptive computational algorithms converge to deliver unparalleled single-cell sensitivity and sub-cellular spatial precision, a potentially transformative tool for biomedical research and diagnostics.

About two decades ago, single-molecule fluorescence in situ hybridization (smFISH) paved the way for spatially resolved transcriptomics by enabling visualization of individual RNA molecules within cellular contexts^38^. However, smFISH was initially limited by low multiplexing capacity, restricting its ability to capture more than a handful of transcripts per experiment. Further technologies made notable advances, expanding multiplexing capabilities and providing finer spatial resolution. Despite these improvements, achieving a commercially viable whole transcriptome capability remained out of reach.

Parallel to FISH advancements, single-cell RNA sequencing (scRNA-seq) emerged as a powerful tool to profile transcriptomes at single-cell resolution. scRNA-seq revealed the complexity of cellular diversity across tissues, playing a pivotal role in cell atlasing initiatives and driving discoveries in cancer, immunology, neuroscience, and developmental biology. However, traditional scRNA-seq requires dissociating cells from their native environments, which can introduce biases, lead to loss of delicate or tightly packed cells, and sacrifice spatial information crucial for understanding tissue architecture and cellular interactions^7^. Spatial transcriptomics profiling tools, such as GeoMx DSP, HDST, Slide-seq and Stereo-seq, addressed some of these limitations by preserving spatial context, but they lacked single and sub-cellular spatial resolution and struggled to localize individual transcripts in three dimensions^39^.

The CosMx WTx assay overcomes these historical limitations by achieving whole-transcriptome coverage *in situ*, by coupling robust assay chemistry with leading-edge hardware and software. For the first time, imaging-based spatial transcriptomics has reached the level of depth and coverage previously achievable only with sequencing-based methods, without sacrificing spatial resolution or single-cell specificity. The key features and applications for CosMx WTx are summarized below.

### High-Resolution, High-Throughput Data Output

CosMx WTx leverages cutting-edge wide field custom optics (over 0.25 mm^2^ FOV at 1.1NA) and low volume microfluidics to economically process high-density samples, enabling the profiling of millions of cells with ∼ 3 cm^2^ imaging area per slide. With an unprecedented localization accuracy of ≤ 50 nm in the xy-plane and ∼600 nm in the z-plane, CosMx WTx captures spatial details that can drive new insights into cellular and sub-cellular organization, tissue architecture, and disease pathology. This level of spatial resolution is particularly valuable for exploring complex tissue microenvironments where cellular interactions play critical roles, such as in tumors and immune niches.

### Adaptive Decoding Algorithm for Enhanced Data Quality

One of the limitations of previous spatial transcriptomics approaches has been the difficulty of distinguishing true transcript signals from background noise, especially in complex tissue samples. The CosMx WTx assay employs a state-of-the-art adaptive decoding algorithm, which dynamically adjusts to maximize transcript capture while minimizing non-specific signals. This technological advancement greatly enhances data quality, providing higher confidence in transcript counts and enabling more precise single-cell analyses.

### Order-of-Magnitude Higher Cell and Transcript Recovery Rate in FFPE Samples

Formalin-fixed, paraffin-embedded (FFPE) tissue samples present unique challenges due to crosslinking and RNA degradation. The CosMx WTx assay demonstrates remarkable sensitivity and specificity in FFPE tissues, achieving one order of magnitude higher cell and transcript recovery rate than standard scRNA-seq from the same tissue volume (e.g., a 5 µm section). By avoiding the cell dissociation process required for scRNA-seq, CosMx WTx also mitigates biases associated with selective cell loss, allowing for a more accurate representation of cellular composition within intact tissue structures^7^.

### Broad Applicability Across Tissue Types

Unlike many spatial transcriptomics tools, which may be optimized for specific tissue types such as brain, CosMx WTx has demonstrated broad applicability across a range of tissue types. The ability to utilize archival FFPE tissue sections ensures that the vast number of key health-care related tissues can be measured in order to significantly impact our understanding of health and disease. This flexibility is critical for translating the technology into diverse research and clinical applications, from oncology and neuroscience to developmental biology and regenerative medicine.

### High Dynamic Range in Transcript Detection

CosMx WTx’s ability to detect over 50,000 transcripts per cell in FFPE tissue (see Table 1, maximum transcripts per cell) highlights that the chemistry is capable of detecting transcripts at this high level. This level of transcriptome coverage within individual cells allows for detailed pathway analysis, enhanced identification of rare or subtle cell states, and exploration of gene expression at a granular level. Such depth is unprecedented for an imaging-based platform.

### Comprehensive Coverage for Spatially Resolved Pathways and ligand-receptor Analysis

The whole-transcriptome capability of CosMx WTx facilitates complex pathway analyses and ligand-receptor interaction studies within a spatial context, offering new dimensions of information that were previously inaccessible. By preserving the spatial organization of tissues, CosMx WTx allows researchers to link gene expression patterns to cellular positioning, tissue structure, and intercellular communication, providing richer insights into disease mechanisms and potential therapeutic targets.

### Cloud-Based Data Storage, Processing, and Sharing

In line with the needs of modern, data-intensive spatial biology, the CosMx WTx assay is supported by an integrated cloud platform (AtoMx Spatial Informatics Platform) for data storage, processing, and sharing. This cloud-based solution streamlines data management, ensuring secure and scalable storage, rapid processing, and easy sharing with collaborators. By simplifying data handling, the AtoMx SIP accelerates research workflows and facilitates collaborative efforts, making high-throughput spatial transcriptomics accessible to a broader audience.

## Implications and Future Directions

The CosMx WTx assay’s ability to combine spatial precision with whole-transcriptome coverage has broad implications for biomedical research. For the first time, researchers can profile tens of thousands of genes across millions of cells while preserving each transcript’s spatial context, empowering studies that demand both profiling depth and spatial resolution. This breakthrough is particularly relevant for fields that rely on understanding cell-to-cell interactions, such as cancer biology, immunology, and developmental studies. In cancer research, for instance, CosMx WTx enables mapping of tumor microenvironments, tracking immune infiltration patterns, and understanding heterogeneity at the single-cell level, which could lead to more targeted and effective therapies.

The capability to resolve single transcripts in three dimensions also opens new avenues for studying cellular structures and their functional implications within tissues. Moving forward, CosMx WTx could potentially be extended to include multi-omic profiling, incorporating protein and other biomolecular markers to achieve a more comprehensive view of cellular states and tissue architecture.

In conclusion, the CosMx WTx assay redefines the boundaries of single-cell and spatial transcriptomics, addressing longstanding limitations of existing technologies. By merging high-resolution imaging with whole-transcriptome coverage, CosMx WTx offers a powerful and versatile tool for spatially resolved molecular profiling. We anticipate that this platform will not only enhance current research but also drive new discoveries and translational applications, establishing a new standard for single-cell and spatial analysis.

## Methods

### Data Availability

Data will be made publicly available as part of a formal peer-review process. Code to reproduce figures can be found at https://github.com/Nanostring-Biostats/WTX_CosMx_SMI.

### Cell culture and slide preparation

Adherent cell lines HEK293T, SH-SY5Y, and U2OS were purchased from ATCC. The cells were cultured in 0.2 µm vacuum filtered DMEM (ATCC), DMEM/F12 (Gibco), and McCoy’s 5A (ATCC) media with 10% fetal bovine serum (Gibco) and 1% Penicillin-Streptomycin (Gibco) respectively. All cell lines were maintained in a humid incubator supplemented with 5% CO_2_ at 37°C.

8-well slide chambers (Nunc) were applied to UV sterilized, poly-L-lysine coated microscope slides (Electron Microscopy Sciences). Confluent cell cultures were trypsinized and resuspended in 200-500 µL growth media such that 50,000-100,000 cells could be applied to the four middle wells (within the CosMx scanning area) of the slide chamber. Slides were gently agitated to distribute cells evenly within each well and put back to incubator for 6-8 hours.

Cells were then inspected under bright field microscope to ensure proper adhesion. The wells were gently washed with 1xPBS, and ice cold 4% PFA (Thermo Scientific) was applied for 30 min. Cells were washed thrice post-fixation with 1xPBS (ThermoFisher). The culture chamber was then removed, and the samples were incubated with 70% ethanol (Pharmco) for 2 min. Slides were transferred to fresh 70% ethanol for another 2 hours or stored at −20°C for up to one month.

### CosMx WTX assay on cultured cells

Slides with cultured cells were incubated in 50% ethanol and 1xPBS for 5 min each to rehydrate the sample. Cells were subsequently permeabilized in 1xPBS with 0.1% Tween-20 (Teknova) for 10 min and washed in 1xPBS twice for 1min each. Incubation frames (Bio-Rad) were then applied to the slide and the sample was digested using 5 µg/mL-1 endoproteinase Glu-C (New England Biolabs) at 40°C for 15 min. Slides were rinsed with DEPC water and incubated in a light-protected chamber with 0.0003% fiducials (Bangs Laboratories) diluted in 2xSSC with 0.1% Tween-20 for 5 min. Samples were washed in 1x PBS for 1 min and fixed in 10% neutral buffered formalin for 1 min.

Slides were immediately quenched with Tris-glycine buffer (0.1 M Tris, 0.1 M Glycine) twice for 5 min each and then 1x PBS for 5 min. Reactive amines were blocked by incubating the sample with 100 mM NHS-acetate (ThermoFisher) in reaction buffer (0.1 M NaP, 0.1% Tween, pH 8.0 in DEPC water). Samples were washed with 2x SSC for 5 min.

WTx ISH probes were denatured at 95°C for 2 min and snap-cooled on ice. The final probe mix (consisting of 0.5nM denatured probes, 40% formamide, 2.5% dextran sulfate, 0.2% BSA, 100 μg/mL salmon sperm DNA, 2xSSC, 0.2 U/µL SuperaseIn (Thermo Fisher)) was applied to the samples. The samples were placed in a hybridization tray with dampened Kimwipe overnight at 37°C. The samples underwent two 25 min stringent wash with 50% formamide in 2xSSC at 37°C to remove unbound probes. Slides were then washed with 2xSSC twice for 2 min. The DAPI nuclear stain was applied to the samples at 1 µg/mL for 15 min at room temperature. Following nuclear staining samples were washed in 1xPBS for five minutes, the segmentation antibody mix (Bruker Spatial Biology) was applied at 0.5x diluted in RNA Blocking Buffer (Bruker Spatial Biology) and incubated with the sample at room temperature for 1 hour. Slides were then washed three times in 1xPBS for 5 min each, and a flow cell (Bruker Spatial Biology) was applied prior to instrument run.

### SMI Instrument

SMI utilizes standard sample preparation methods typical for FISH or immunohistochemistry on FFPE tissue sections. In addition, fluorescent bead-based fiducials are added to the tissue which is then fixed to provide an optical reference for cyclic image acquisition and registration. For sample-processing details, see the “RNA assay FFPE tissue prep by manual method” section (below). Following hybridization of ISH probes or antibody incubation, slides were washed and the coverslip with a defined height (70 μm) spacer was applied to the slide. The slide plus coverslip constitutes the flow cell, which was placed within a fluidic manifold on the SMI instrument for readout and morphological imaging. The *in situ* chemistry and imaging analyses were performed on a commercial-grade CosMx instrument with a custom large FOV, high numerical aperture, and low aberration objective (and associated optics) that permit more than 3,000 cells to be imaged per FOV from a flow cell assembled onto a standard glass slide. The system can run up to 4 flow cells simultaneously and can interleave fluidics and optical-imaging operations to maximize throughput. The optical system has an epi-fluorescent configuration that is based on a custom water objective with a numerical aperture (NA) of 1.1 and a magnification of 22.8X. The FOV size was customized to 512 μm x 512 μm. Illumination is widefield with a mix of lasers and LEDs that allow imaging of Alexa Fluor-488, Atto 532, Dyomics Dy-605, and Alexa Fluor-647 as well as cleaving of photolabile-dye components. The camera used is a Lucid ATX204S 10GigE based on the IMX531 Sony CMOS sensor, and the sampling at the image plane is 120 nm/pixel.

An XY stage moves the flow cell above the objective lens, and a Z-axis motor moves the objective lens. The fluidic system uses a custom interface to draw reagents through a flow cell using a syringe pump. Reagent selection is controlled by a shear valve (Idex Health & Science). A flow sensor between the flow cell and the syringe pump was used for flow rate feedback (Sensirion AG). The fluidic interface includes a flat aluminum plate that is in direct contact with the flow cell. The temperature of this metal plate was controlled to regulate the reporter hybridization temperature at 28°C. The enclosure around the opto-mechanics and flow cell holder inside the instrument was also maintained at a constant temperature using a separate thermoelectric cooler.

### Secondary analysis and decoding

A decoding algorithm is used to translate reporter-binding events (RBE) (characterized by x,y,z location of the imaging-barcode colors into transcripts (characterized by x,y,z location and gene target identity). Multiple RBE in close spatial proximity which correspond to a gene-specific barcode provide evidence for a gene transcript call at a consensus location of the corresponding RBE, and negative signals can be used to calibrate the accuracy of the decoding algorithm and the level of nonspecific background on the tissue (**Figure 2A-B**). The assay contains two types of negative signals: negative controls and system controls. Negative controls are described above and measure the intrinsic affinity of any nucleic acid to the surface, as the target for the probes is not in the sample. System controls measure decoding error, as no probe containing a system control barcode is present in the sample, but counts can be assigned to that barcode during target decoding. An important goal of the decoding algorithm is to use the built-in controls to generate quality scores for each transcript which can be used to optimize specificity of the decoding process and filter out transcript calls which behave similarly to system controls (often called “false codes” in the literature) or negative probe controls. To boost sensitivity of the decoding process, we allow for partial barcode matching, such that a transcript may be called when only 3 of the 4 on-bits for that gene’s barcode are observed. This can be justified by the barcode design (HW4, HD >=4), where each gene’s unique barcode with 4 on-bits is composed of 4 unique target-specific combinations of 3 on-bit barcodes. Transcript level quality scores account for all relevant metrics, including the completeness of the barcode match.

The decoding algorithm works in 4 main stages. It should be emphasized that, due to the sheer density of data and the computational demands on decoding high-plex signals, the decoding operation is performed in the cloud (in the AtoMx Spatial Informatics Platform). The first stage of the algorithm identifies all potential transcripts where at least 3 out of 4 on-bits in a gene’s barcode are observed among RBE within a 1.1 pixel radius of the transcript centroid, and calculates a number of quality metrics indicative of the likelihood that the potential transcript is ‘real’, or behaves more like a system control code (random grouping of RBE), or negative control (non-specific background).

Metrics computed at this stage are based on crowdedness, spatial distribution, and barcode likelihood in the local neighborhood of RBE surrounding the call.

The second stage of the algorithm directly uses system control transcript calls as built-in controls, fitting a machine learning model predicting whether each potential transcript call is “not a system control code”, using quality metrics obtained in stage 1 as covariates^40,41^. The output prediction is a system control quality score which ranges between 0 and 1, with higher scores indicating confident transcript calls which do not have similar quality characteristics as the built-in system controls (**Figure 2C**).

In the third step of the algorithm, transcripts are ranked by their system control quality score from stage 2, and then filtered until the total proportion of system controls across all potential remaining transcripts in a FOV is less than 5 percent. Duplicate transcript calls are removed such that two transcripts of the same gene are no closer than 0.75 pixels from each other. Several additional metrics related to transcript quality are then calculated, using the remaining potential transcripts, based on the spatial distribution of the remaining transcript calls, including metrics related to local background levels. The fourth stage of the algorithm works to create a “negative probe quality score”, using a similar process as that described in the second stage, but using a different set of metrics to predict whether each transcript is “not a negative probe” (**Figure 2C**). Following the fourth stage, each transcript has a system control quality score and a negative probe quality score, which can be used to assess overall voted-call confidence at the transcript level. A subset of high confidence transcript calls based on full barcode matches is used to calibrate the full set of voted calls and determine a two-dimensional threshold for transcript inclusion to carry forward in a tertiary analysis. Transcripts are accepted for inclusion in the dataset based on both negative probe and system control quality scores (**Figure 2D**).

To infer the precision of estimated transcript call locations, we used the distances of the transcript locations to their assigned RBE’s to generate 95% confidence intervals (CI)’s for each transcript’s location in x and y dimensions. Using all transcript from the first 10 FOV of the FFPE colorectal cancer dataset (14,578,769 transcripts), we found the average CI width to be < 40 nm, with a standard deviation of ∼12 nm (**Supplementary Figure 1**).

### Data analysis for cell-based WTx assay

For the reproducibility test, two different batches of HEK293T cells were used. Correlation analysis with bulk RNA-seq was performed by using the public dataset retrieved from NCBI GEO (GSM3734788). Downstream analyses, including data normalization, transformation, dimension reduction, unsupervised clustering and differential expression used the Giotto v4.0 package in R^42^.

### Benchmarking CosMx WTx against scRNA-seq in FFPE colon cancer

scRNA-seq for FFPE colon cancer tissues followed the standard best-practice Chromium Flex assay protocol and performed by an exclusive 10x Genomics Clinical Research Organization – Azenta Life Sciences. In brief, two 25 µm tissue curls were subjected to dissociation using the Miltenyi FFPE tissue dissociation kit, and 10,000 cells were then used for barcoding and library preparation, resulting in ∼63% cell recovery rate. The primary scRNA-seq data was processed with CellRanger software, with ∼79.64% reads confidently mapped to cells (median 1,151 UMIs and 774 genes per cell). Downstream analysis was performed with Seurat v5.0 package^26^. In general, cells with >5% mitochondrial counts or with total counts at the top 5% of population were filtered out. Then the data matrix went through log1P and z-score transformation, PCA and UMAP dimension reduction. Leiden clustering was performed to group cells, followed by cell type annotation based on differential gene expressions. Due to the limited number of cells in scRNA-seq, only a coarse annotation was performed, where seven primary cell types were identified and used for comparison. For the CosMx WTx colon cancer dataset, 400 FOVs (single-FOV dimension 0.51 mm x 0.51 mm) were placed and yielded about half million segmented cells. Cells with less than 200 transcript counts or 50 gene features or larger than 5x geometric mean cell size were filtered out. Data processing followed the same principle as scRNA-seq, with the matched seven primary cell types annotated. However, with a significantly larger number of cells the CosMx dataset enabled much higher granularity and finer cell types/subtypes to be identified (implemented in the following analysis). To test the cell composition difference between datasets, ∼6,000 cells were randomly sampled from the CosMx dataset for 100 times, and the average cell composition was presented in Figure 5D, while the cell composition of the entire CosMx dataset was presented in Figure 5E.

### Design of the Whole Transcriptome Atlas probes

The NCBI RefSeq reference transcriptome for human (T2T-CHM13v2.0) was used for design of human WTx, as accessed in May 2023. The genes targeted included all protein-coding genes with a few exceptions, including the high expressing genes below and all mitochondrially-encoded protein-coding genes. Protein coding genes were determined based on accessing the HGNC data July 6, 2023. The 10 high expressing genes excluded from the panel are as follows: *ACTB*, *ACTG1*, *EEF1A1*, *EEF2*, *FTL*, *GAPDH*, *PSAP*, *RPL3*, *TPT1*, and *UBC*. *CARD8* and *CPAMD8* were excluded from these experiments due to spuriously high signal later corrected by probe redesign.

In the probe design process, all possible contiguous 35–50 nucleotide sequence windows for each mRNA target were evaluated. The pool of candidates was first filtered for intrinsic characteristics including melting temperature (Tm), GC content, secondary structure, and runs of polynucleotides. Probes satisfying these parameters were further screened for homology with the full transcriptome of the parent organism using the Basic Local Alignment Search Tool (BLAST)^43^. Preference was given to probes with absence of homology with off-target genes, probes covering known protein-coding transcripts, and maximizing the coverage of the isoform repertoire. Targeting of a transcript was judged based on ≥95% sequence identity to the probe target. Previous work has found that selecting probes that are 95%–100% identical to the intended target and filtering out probes that are ≥75%–85% in homology and that possess ≥15– 17 Maximum Contiguous Bases (MCB) confer excellent specificity to the intended target^44–46^. Final panel candidates were further screened for intermolecular interactions with other probes in the candidate pool including potential probe–probe hybridization as well as minimizing common sequences between probes. Finally, as T2T is a newer genome build, we ensured that all probes targeting genes common between T2T and GRCh38 hit transcripts aligning to both builds.

Negative control probes were designed against synthetic sequences from the External RNA Controls Consortium (ERCC) set^17^. Lack of similarity to any known transcripts was confirmed by BLAST comparison to each transcriptome for all selected negative sequences. Negative control probes were designed to have similar GC and Tm properties as target probes and are subject to the same intermolecular interaction screening.

### Transcript Length Analysis and Karyotype Plot

For the transcript length histogram in Figure 1A and the density map in Figure 1B, all targeted accessions are based on BLAST of probe sequences to human NCBI RefSeq sequences present in our internal database July 2024, updated from RefSeq on June 19, 2024. Whether an accession is considered targeted is based on global percent identity of 95% or greater. The histogram was made using the Python Seaborn histogram function, with all values >15,000 grouped into a single bin. The karyotype plot was made using RIdeogram^47^. A custom karyotype was generated based on the newest human genome, T2T CHM13v2.0/hs1. Data was accessed July 23, 2024 from the T2T GitHub page^48^. RefSeq accession positions in the T2T genome build were accessed from the UCSC table browser on July 23, 2024^49^. The gene density plot was made by normalizing the number of accessions targeted by the panel per 10^6^ bases (1 Mb).

### Application of the CosMx WTx Assay to Colon Cancer

Decoded data from the colon adenocarcinoma FFPE sample went through the following workflow: quality control (QC), pre-processing, cell typing, tertiary analysis of pathways and ligand receptors.

For QC, a simple cellular filter was applied to the dataset where cells with greater than 400 total counts were retained (84% of cells). Since it is unexpected that every cell produces high expression of every gene, low-expressing targets were removed. Specifically, targets whose average expression was above three standard deviations of the mean expression of the negative controls (0.031) were retained. This procedure reduced the feature count to 13,918 out of 18,935 targets (73.5%). The filtered dataset was then converted to annData format and normalized, log1p-transformed, and scaled with SCANPY^50^. Dimensional reduction was performed using the first 50 principal components, 50 nearest neighbors, and UMAP parameters min_dist=0.2 and spread=0.8.

A two-pronged approach was used to classify cell types. The first approach identified tumor cells and the second approach classified the various non-tumor cell types. For tumor cell determination, all cells were clustered using Leiden Clustering with a resolution of 0.9. Based on marker gene expression, four out of 20 clusters were associated with canonical tumor markers (*e.g.*, *CEACAM5*). Second, the non-cancerous Intestines CellTypist database was used to create a matrix of average expression of 36 cell types^51,52^. The fully supervised InSituType algorithm^53^ then classified all cells using raw counts and the mean expression of negative probes as background. The two approaches were then combined where cells belonging to one of the four tumor clusters from Leiden approach were classified as tumor and the remaining cells were classified from InSituType. Since the primary interest for this study was on local pathway scores and local LR interactions within the tumor nest, the 40 categories were combined into 25 broader cell type classifications.

The PROGENy Database was used with decoupleR^54^ to quantify the local pathway enrichment scores across the colon sample^27^. This was done by computing the spatial weights using a Gaussian kernel in the package LIANA+. For this procedure, a bandwidth of 150 and a cutoff of 0.1 was used. The multivariate linear model from decoupleR was used estimate pathway enrichment throughout the tissue and globally.

LIANA+ and decoupleR were also used to identify spatially co-expressed LR pairings within the tumor nest. This was done using the bivariate method of LIANA+ with 100 permutations. The LR pairs with the greatest local score standard deviation were then identified to quantify intra-tumoral cellular communication variation.

