## Supplementary Figures and Supplementary Table 1 for "Sub-cellular Imaging of the Entire Protein-Coding Human Transcriptome (18933-plex) on FFPE Tissue Using Spatial Molecular Imaging"

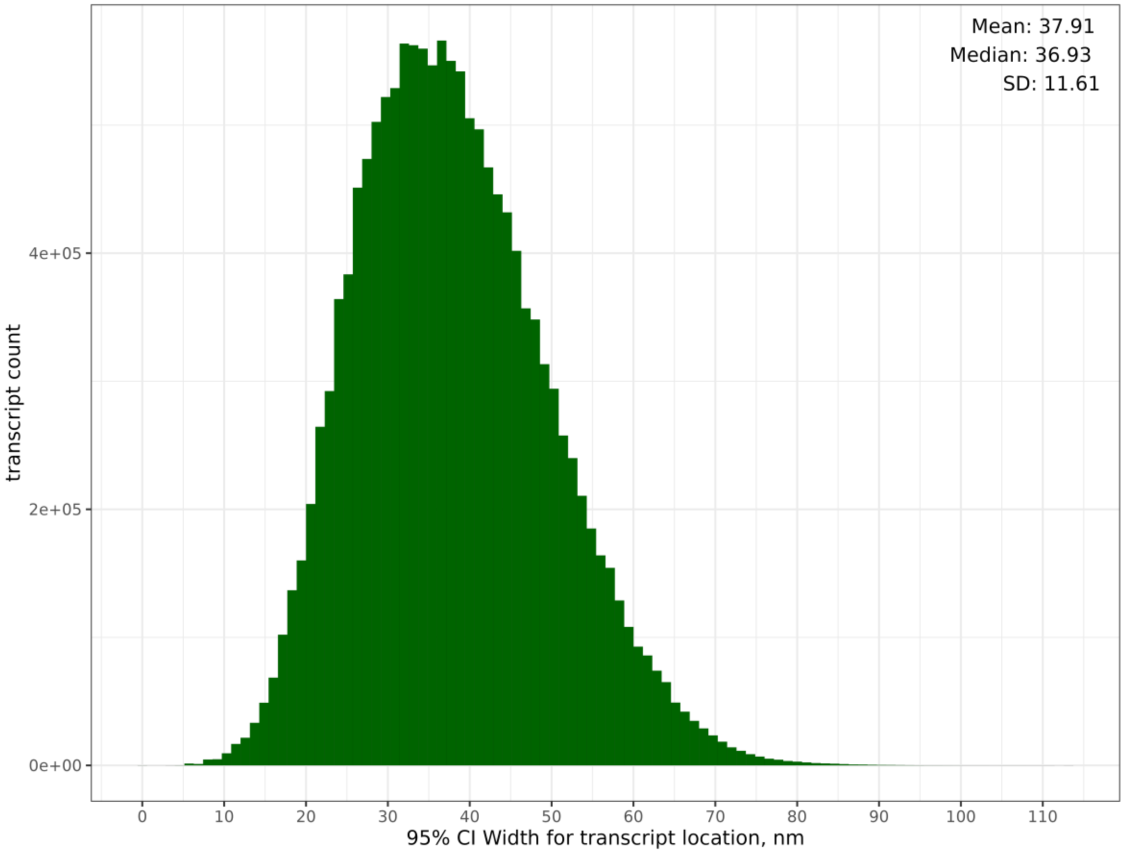


Histogram of confidence interval (CI) widths in nm for transcript locations, calculated using the distances of reporter binding events from the transcript call location in X and Y planes. Histogram based on first 10 FOVs of WTx Colon Cancer dataset (14,578,769 transcripts)

**Supplementary Figure 1**

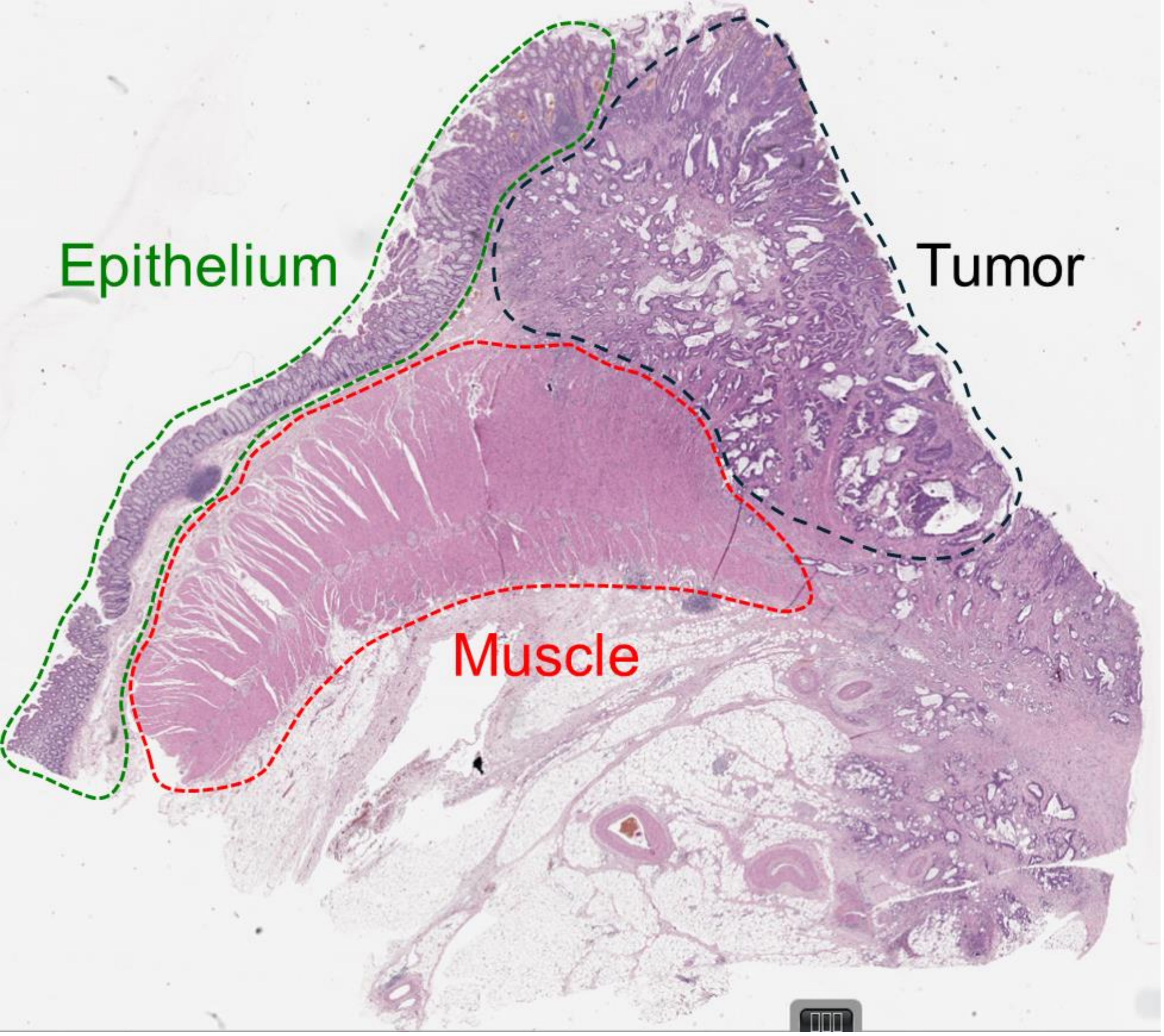


H&E image of tissue sample prepared for scRNA-seq. Areas containing key cell types are indicated.

**Supplementary Figure 2**

| Performance metrics | Chromium | CosMx |
|---|---|---|
| Total number of genes | 18,082 | 18,935 |
| Input sample | 10,000 cells | 400 FOVs |
| Output number of cells | 6,275 | 493,929 |
| Median counts/cell | 1,151 | 967 |
| Median features/cell* | 774 | 627 |
| NegativeProbe/plex/cell | NA | 0.01457 |
| FalseCode/plex/cell | NA | 0.00539 |
| Genes over background | NA | 13,918 |

| Primary cell type | Chromium # | Chromium % | CosMx # | CosMx % |
|---|---|---|---|---|
| Smooth muscle | 278 | 5% | 71,249 | 19% |
| Stroma cell | 1,234 | 21% | 96,517 | 26% |
| Endothelial | 286 | 5% | 6,650 | 2% |
| Goblet/Epithelial | 180 | 3% | 52,239 | 14% |
| Lymphocyte | 556 | 9% | 22,904 | 6% |
| Macrophage | 1,042 | 18% | 28,971 | 8% |
| Tumor | 2,378 | 40% | 96,272 | 26% |
| **Total** | 5,954 | 100% | 374,802 | 100% |

**Supplementary Table 1**
